# Structural basis for pore blockade of the human cardiac sodium channel Na_v_1.5 by tetrodotoxin and quinidine

**DOI:** 10.1101/2019.12.30.890681

**Authors:** Zhangqiang Li, Xueqin Jin, Gaoxingyu Huang, Kun Wu, Jianlin Lei, Xiaojing Pan, Nieng Yan

## Abstract

Among the nine subtypes of human voltage-gated Na^+^ (Na_v_) channels, tetrodotoxin (TTX)-resistant Na_v_1.5 is the primary one in heart. More than 400 point mutations have been identified in Na_v_1.5 that are associated with cardiac disorders exemplified by type 3 long QT syndrome and Brugada syndrome, making Na_v_1.5 a common target for class I antiarrythmic drugs. Here we present the cryo-EM structures of Na_v_1.5 bound to quinidine and TTX with resolutions at 3.2 Å and 3.3 Å, respectively, for the pore domain. The structures allow mapping of 278 disease-related mutations in the resolved region, establishing the framework for mechanistic investigation of these pathogenic mutations. Quinidine is positioned right below the selectivity filter and coordinated by residues from repeats I, III, and IV. Pore blockade is achieved through both direct obstruction of the ion permeation path and induced rotation of a Tyr residue that leads to intracellular gate closure. Structural comparison of Na_v_1.5 with our previously reported Na_v_1.2/1.4/1.7 reveals the molecular determinant for the different susceptibility to TTX by TTX-resistant and -sensitive Na_v_ subtypes, and highlights the functional significance of the underexplored and least conserved extracellular loops.

## Introduction

Each heartbeat is a result of sophisticated cooperation of multiple signaling events, including transmission of electrical impulses, conversion of the membrane electrical signals to the intracellular chemical signals, and eventually the contraction of cardiomyocytes (1, 2). Among these, the initiation of an action potential is mainly mediated by the cardiac subtype of voltage-gated sodium (Na_v_) channels, Na_v_1.5, which is insensitive to tetrodotoxin (TTX) and encoded by the gene *SCN5A (3)*. Due to its essential physiological function, it is not surprising that among the over 1000 identified disease mutations known in the nine human Na_v_ channels, more than 400 have been found in Na_v_1.5 (Tables S1-S3) (4).

Gain or loss of function (GOF or LOF) of Na_v_1.5 can both be pathogenic (5-11). LOF mutations, which may result in reduced Na^+^ peak current *I*_*Na*_ or prolonged inactivation, are associated with Brugada syndrome, sick sinus syndrome, and cardiac conduction defect. GOF mutations can lead to type 3 Long QT (LQT3) syndrome, sudden infant death syndrome, and stillbirth. A common arrhythmia, atrial fibrillation, can be associated with either GOF or LOF.

Na_v_1.5, on one hand, is a primary target for class I antiarrythmic drugs, which are Na_v_ blockers and further classified to Ia, Ib and Ic (12). On the other hand, Na_v_1.5, similar to the cardiac potassium channel hERG, has to be avoided for general drug development due to potential off-target activity (13, 14). Therefore, a detailed structure of human Na_v_1.5 is required to understand its pathophysiology and to facilitate drug discovery.

In the past three years, we have reported the cryo-EM structures, at resolutions ranging between 2.6-4.0 Å, of Na_v_ channels from insect (Na_v_PaS), electric eel (EeNa_v_1.4), and finally human, including Na_v_1.2, Na_v_1.4, and Na_v_1.7 that function respectively in the central nervous system, skeletal muscle, and sensory neurons (15-20). Apart from the first structure of a eukaryotic Na_v_ channel, Na_v_PaS, whose pore domain has no fenestration and VSDs are only partially “up”, all other Na_v_ channels exhibit similar conformations with voltage-sensing domains (VSDs) in the depolarized “up” states and the fast inactivation motif plugging into a structurally-revealed receptor cavity between the S6 pore helices and the S4-S5 restriction helices in repeats III and IV. Because of the high sequence similarity, the structures of these Na_v_ channels in complex with distinct auxiliary subunits and small molecule or peptidic toxins, elucidate the folding principle of the single chain voltage-gated ion channels, the details of the Na^+^ permeation path through the Asp/Glu/Lys/Ala (DEKA)-constituted selectivity filter (SF), the mechanism of fast inactivation, and the molecular basis for specific recognition and mechanism of action of various modulators. These structures also provide the molecular template to study hundreds of disease mutations.

On the basis of the previous progress, our incentive for structural investigation of Na_v_1.5 is to unveil its interplay with representative ligands and to identify distinct features that may facilitate development of specific therapeutics targeting different Na_v_ subtypes. Here we present the structures of full-length human Na_v_1.5 in the presence of TTX or quinidine, a class Ia antiarrythmic drug, determined using single-particle cryo-EM. To avoid redundancy with previous structural illustrations, we focus our analysis on the unique features of Na_v_1.5.

## Results

### Cryo-EM analysis of human Na_v_1.5

Full-length human Na_v_1.5 was transiently co-expressed with the β1 subunit in HEK293F cells using plasmids and purified through tandem affinity column and size exclusion chromatography (Figure 1A). We chose β1 to be co-expressed with Na_v_1.5, because TTX-resistant Na^+^ current was elevated when Na_v_1.5 was co-expressed with β1, although there was little effect on the activation and inactivation curves (Figure 1B, Figure S1, Table S4). This observation is consistent with the previous report of increased expression level of Na_v_1.5 in the presence of the β1 subunit (21).

**Figure 1.**
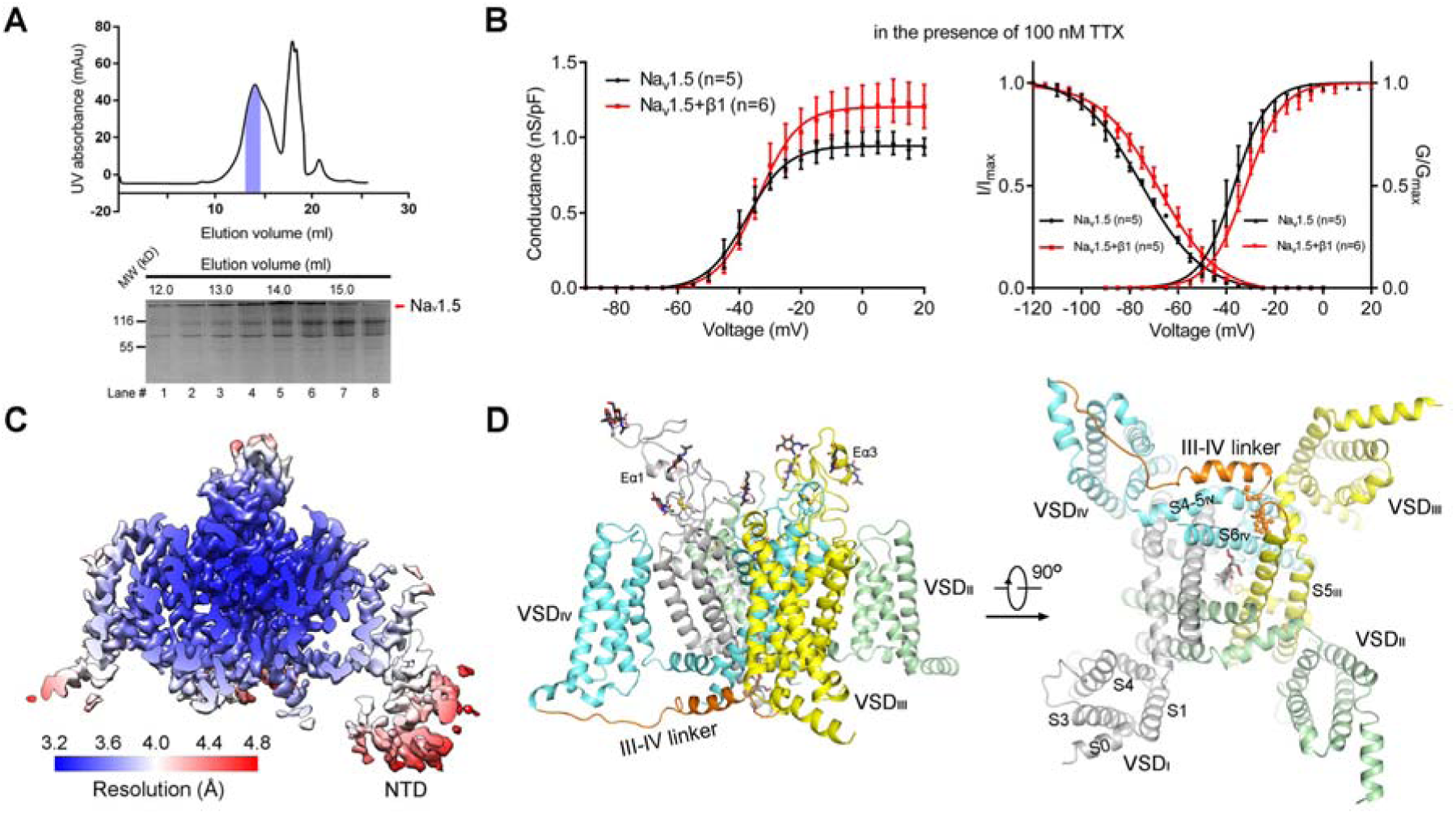
Structural determination of full-length human Na_v_1.5. (**A**) Size-exclusion chromatography (SEC) purification of human Na_v_1.5 co-expressed with the β1 subunit in HEK293F cells. Shown here is a representative chromatogram of gel filtration purification. The indicated fractions were concentrated for cryo sample preparation. (**B**) Co-expression with the β1 subunit increases Na^+^ current without significant shift of the electrophysiological property of Na_v_1.5. All the recordings were performed in the presence of 100 nM tetrodotoxin (TTX). Please refer to Methods for details. *Left*: Conductance densities. *n* values indicate the number of independent cells; mean ± s.e.m. G/G_max_ and I/I_max_ represent normalized conductance and ion current, respectively. *Right*: Voltage-dependent activation and inactivation curves. (**C**) No density corresponding to the β1 subunit is observed. The local resolution map was calculated with Relion 3.0 and presented in Chimera (35). (**D**) Overall structure of human Na_v_1.5. A side and a cytoplasmic view are shown. The structure is color-coded for distinct repeats. The III-IV linker is colored orange and the fast inactivation motif, Ile/Phe/Met (IFM) is shown as ball and sticks on the right. The sugar moieties and a putative GDN molecule are shown as black (left) and grey (right) sticks, respectively.

The freshly purified protein, at concentration of approximately 1 mg/ml, was either incubated with 66 μM tetrodotoxin (TTX) and ∼200 μM untagged calmodulin (CaM), or 1 mM quinidine and 12 μM saxitoxin (STX) for 30 min. The samples, named Na_v_1.5-TC in the presence of TTX and CaM, and Na_v_1.5-QS with quinidine and STX, were applied for cryo-grid preparation, image acquisition, and data processing. EM maps were obtained at overall resolutions of 3.4 Å and 3.3 Å out of 145,139 and 124,954 selected particles, respectively. The resolutions for the central region were respectively improved to 3.3 Å and 3.2 Å after application of a soft mask (Figure 1C, Figures S2-S4, Table S5). The 3D reconstruction maps for Na_v_1.5-TC and Na_v_1.5-QS are similar (Figure S2C). We will focus on one map for overall structural description.

Compared to the 3D reconstructions for human Na_v_1.2 in complex with β2, Na_v_1.4 with β1, and Na_v_1.7 with both β1 and β2, that of Na_v_1.5 apparently lacks the density for any auxiliary subunit although β1 was co-expressed (Figure S2C). Similar to the 3D EM reconstruction of human Ca_v_3.1 (22), the intracellular segments, including CaM, are largely invisible, despite addition of CaM. Instead, the map of Na_v_1.5 contains an extra appendage contiguous with VSD_I_, reminiscent of the amino-terminal domain (NTD) in Na_v_PaS (15, 20) (Figure S2C). However, the resolution for the putative NTD is approximately 4.5-6 Å, insufficient for model building (Figure 1C). The final structural model of Na_v_1.5 contains 1151 residues, including the complete transmembrane domain, the extracellular segments, and the III-IV linker. Nine glycosylation sites were observed and one sugar moiety was assigned to each site (Figure 1D). On the resolved region, 278 disease-related mutations can be mapped, providing the framework for structural-guided mechanistic investigation of these disease variants (Tables S1-S3).

### Pore blockade by quinidine

While TTX was well-resolved in Na_v_1.5-TC, that of STX was only resolved as a blob congesting the entrance to the SF in Na_v_1.5-QS (Figure S4B,C). STX was not modelled. The two structures will hereafter be referred to as Na_v_1.5T and Na_v_1.5Q for complexes with TTX and quinidine, respectively. Unlike Ca_v_1.1 and Ca_v_3.1, which undergo main chain shift of the pore-forming S6 segments when binding to pore blockers such as diltiazem, verapamil, and Z944 (22, 23), the two structures of Na_v_1.5 remain nearly identical the root-mean-square deviation (RMSD) of 0.16 Å over 771 Cα atoms (Figure 2A).

**Figure 2.**
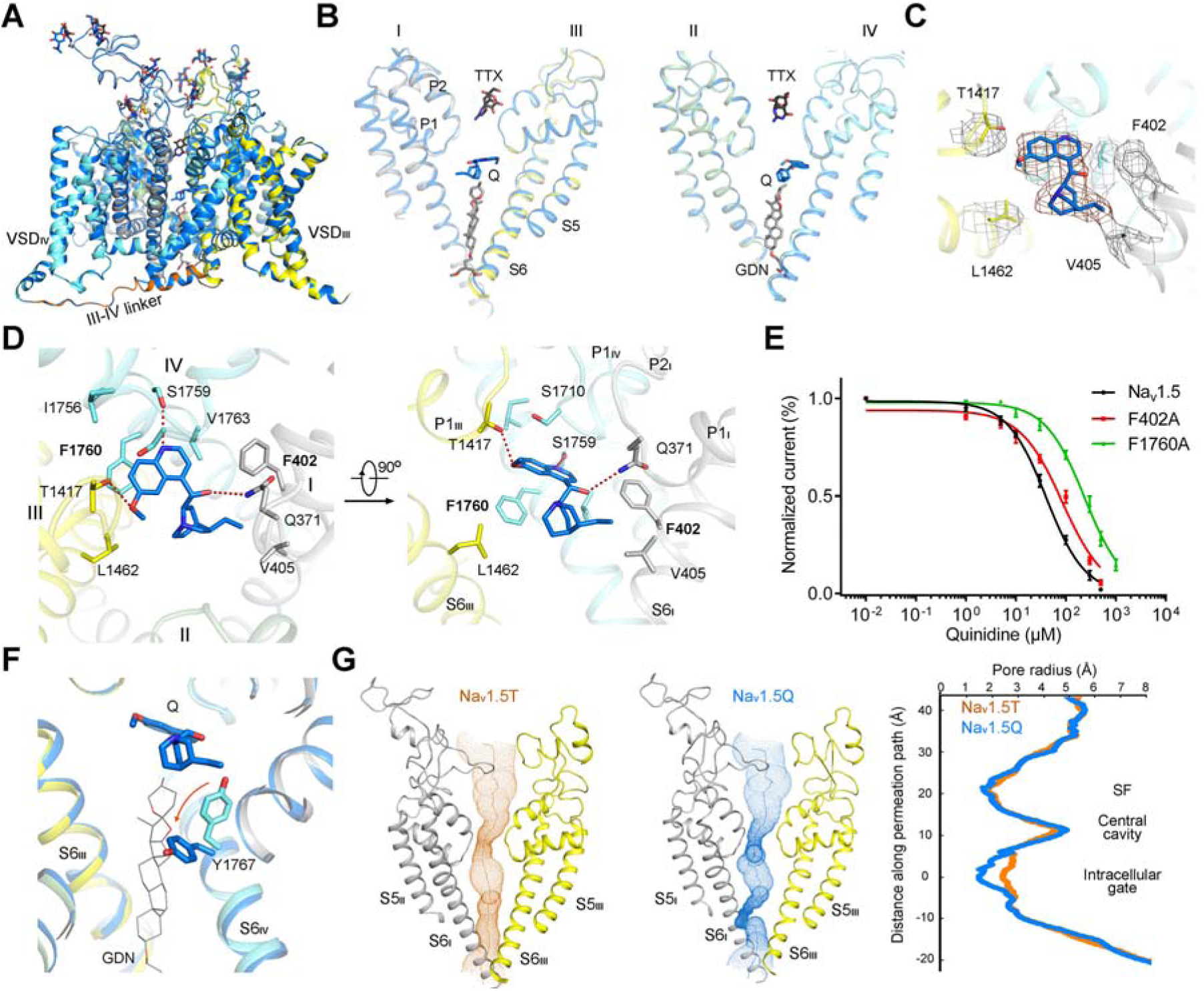
Pore blockade of Na_v_1.5 by quinidine. (**A**) The overall structure of Na_v_1.5 remains unchanged in the presence of TTX or quinidine. The TTX-bound structure is domain colored, and the quinidine-bound one is colored blue. (**B**) Quinidine binds to the central cavity of the pore domain. In the presence of quinidine, GDN no longer penetrates into the cavity through the intracellular gate. (**C**) Local densities for the bound quinidine and surrounding residues. The densities, shown as brown (quinidine) and gray (residues) meshes, are contoured at 5 σ in PyMol. (**D**) Quinidine is mainly coordinated by residues from repeats I, III, and IV. Because of the lack of coordination from repeat II, quinidine is closer to repeat IV, hence off the central axis of the pore domain. Potential hydrogen bonds are indicated by red, dashed lines. (**E**) Validation of the structure-revealed quinidine coordination. Shown here are concentration-response curves for the blockade of Na_v_1.5-WT (black), and variants containing single point mutation F402A (red) or F1760A (green) by quinidine. The Na^+^ currents were measured after perfusion of the drug at indicated concentrations and applying 50-ms pulses at 1 Hz from −120 mV to −20 mV. The IC_50_ values were 40.3 ± 1.1 μM (Na_v_1.5), 84.8 ± 1.3 μM (F402A) and 239.3 ± 1.2 μM (F1760A), respectively, in HEK 293T cells. The sample sizes (*n*) tested from low to high concentrations are: *n* = 11, 7, 9, 6, 8, 4, 5, 5 for Na_v_1.5; *n* = 6, 5, 6, 6, 5, 6, 6, 3 for F402A; and *n* = 7, 4, 5, 6, 6, 6, 7, 7, 3 for F1760A. (**F**) Quinidine binding triggers rotation of Tyr1767, resulting in closure of the intracellular gate. (**G**) The ion permeation path in the pore domain of TTX- and quinidine-bound structures. The permeation path, calculated by HOLE (36), is illustrated by orange and blue dots in the left and right panels. The corresponding pore radii of Na_v_1.5T (orange) and Na_v_1.5Q (blue) are compared in the right panel.

Similar to the structures of EeNa_v_1.4, human Na_v_1.2 and Na_v_1.4, a density that likely belongs to the detergent glycol-diosgenin (GDN) penetrates the intracellular gate of Na_v_1.5T (Figure S4B). However, the GDN density is gone in Na_v_1.5Q because it would clash with the bound quinidine, which is localized in the central cavity of the pore domain (Figure 2B).

The distinctive shape of the quinolone double ring and the quinuclidine “cage” in quinine facilitates ligand fitting into the density (Figure 2C). Consistent with its function as a pore blocker, quinidine is positioned right beneath the SF, with the quinolone plane approximately perpendicular to the central axis and the quinuclidine group further down to the central cavity. Quinidine is coordinated by both polar and hydrophobic residues from repeats I, III, and IV, but distanced from repeat II. Therefore it is off the central axis (Figure 2B,D).

Three polar residues, Gln371, Thr1417, and Ser1759 are likely hydrogen-bonded to the polar groups of quinidine. Gln371 and Thr1417, which are positioned at the bottom of SF in repeats I and III, respectively, appear to hang the blocker below the SF (Figure 2D). Quinidine is surrounded by multiple hydrophobic residues from the S6 segments in the three repeats, among which Phe1760 from S6_III_ appears to play a major role through π-π packing (Figure 2D). Supporting this notion, single point mutation F1760A led to increased IC_50_ from 40.3 ± 1.1 μM to 239.3 ± 1.2 μM, whereas Ala substitution of Phe402, which interacts quinidine through van der Waals contact, only results in marginal increase of IC_50_ to 84.8 ± 1.3 μM (Figure 2E).

Despite the lack of main chain shifts, Tyr1767, the intracellular gating residue on S6_IV_ undergoes local changes. In Na_v_1.5T, Tyr1767 is well-resolved with its aromatic ring pointing up, leaving space for GDN. In Na_v_1.5Q, the density for Tyr1767 becomes much poorer, suggesting its flexibility. Nevertheless, it can no longer keep the original conformation, which would be incompatible with the bound quinidine. The most likely assignment for Tyr1767, indicated from low-thresh hold map, is to flip down to close the intracellular gate, similar to the corresponding residue Tyr1755 in Na_v_1.7 (19) (Figure 2F,G, Figure S4D). Therefore, in addition to directly blocking ion conduction at an upper site, quinidine may also facilitate gate closure.

### Molecular determinant for TTX-resistance

The guanidinium pore blocker TTX has been used as a probe for characterizing Na_v_ channels (24-27). The nine mammalian Na_v_ channels have been classified to TTX-sensitive and TTX-resistant (TTXr), the latter containing Na_v_1.5/1.8/1.9 because they can only be blocked by TTX at micromolar range (27, 28). Sequence comparison and mutagenesis characterizations have pinpointed the determinant to be the replacement of a Tyr or Phe on the pore helix P2_I_ in TTXr by Cys in Na_v_1.5 or Ser in Na_v_1.8/1.9 (27, 29-31). Structures of TTX-bound Na_v_PaS at 2.6 Å and Na_v_1.7 at 3.2 Å show that Tyr is placed close to the guanidinium group (19, 20). We had speculated the hydroxyl group of Tyr to form water-mediated hydrogen bond with TTX, which was challenged by the presence of Phe at this locus in some TTXs Na_v_ channels. We then attributed the interaction between the aromatic ring of Tyr and the positively charged guanidinium group to an unideal π-cation interaction, because their distance is about right, but the electronegative point of the Tyr and the guanidinium group is not optimally aligned for π-cation interaction. Now with the structure of Na_v_1.5T, additional mechanistic insight can be obtained for understanding TTXr and TTXs Na_v_ channels.

The binding site for TTX in Na_v_1.5, mainly constituted by acidic residues at the outer mouth to the SF, is identical to that in Na_v_PaS and Na_v_1.7 (19, 20). We will focus on Na_v_1.7 for comparison (Figure 3A,B). As predicted, Cys373 on P2_I_ in Na_v_1.5, which occupies the same locus for Tyr362 in Na_v_1.7, is not involved in TTX coordination. On repeat III, Na_v_1.7 has two outliers, Thr1409/Ile1410, whose corresponding residues are Met/Asp in other human Na_v_ channels, including Na_v_1.5, and Leu/Gln in Na_v_PaS (20). Consistent with our previous prediction, the backbone amide of Met1422 and the carboxylate of Asp1423 in Na_v_1.5 interact with the C10-OH and C11-OH of TTX, respectively (19) (Figure 3A,B).

**Figure 3.**
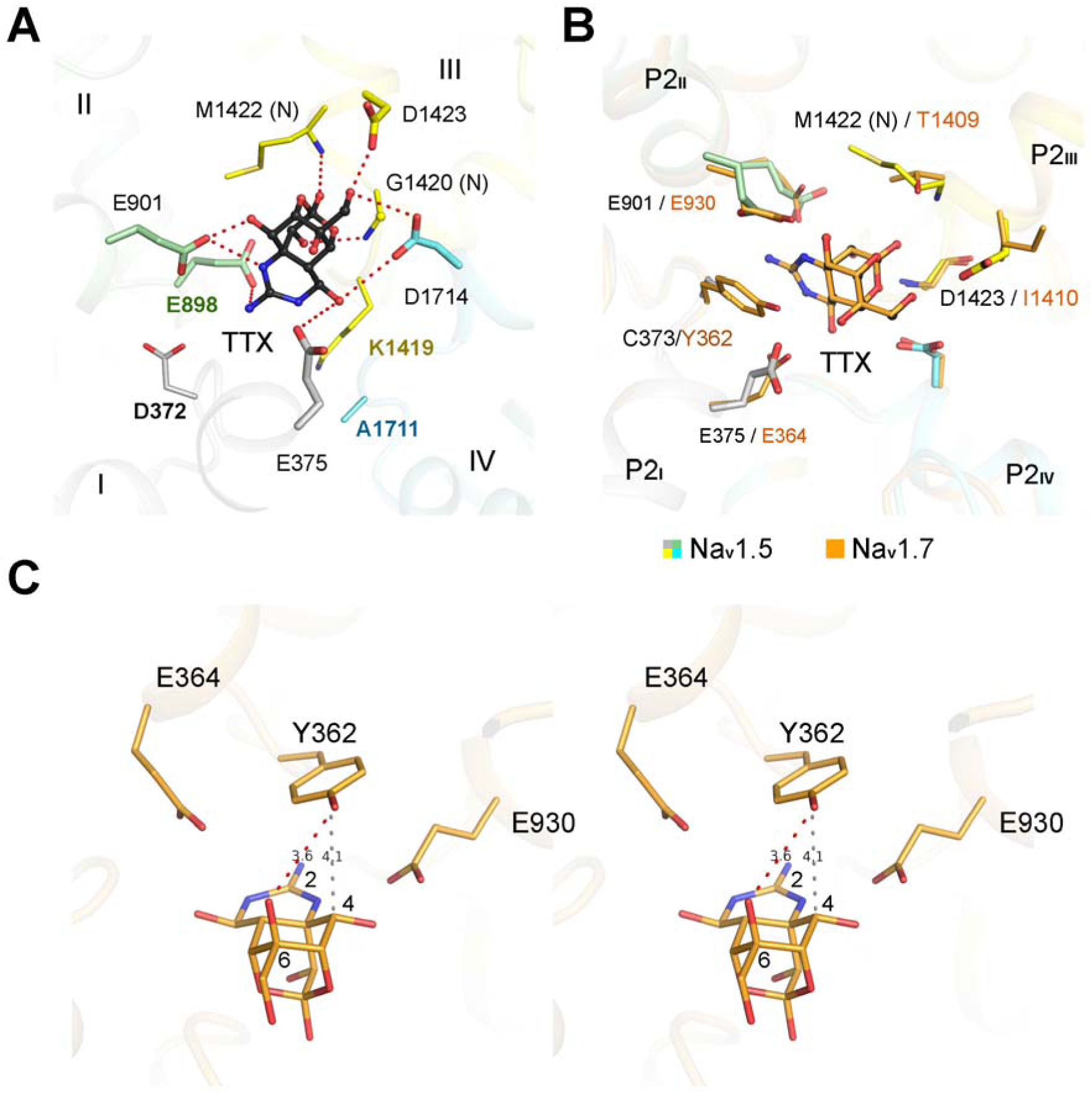
Molecular basis for TTX-resistance by Na_v_1.5. (**A**) Coordination of TTX by Na_v_1.5. An extracellular view of the entrance to the selectivity filter is shown. TTX is shown as black ball and sticks. The DEKA motif residues, Asp372/Glu898/Lys1419/Ala1711, are labeled in colors consistent with their respective domain color. (**B**) Comparison of TTX-coordination by Na_v_1.5 and Na_v_1.7 (PDB code: 6J8J). Thr1409/Ile1410 in Na_v_1.7 represent outliers among human Na_v_ channels, as all others have Met and Asp at these two loci as in Na_v_1.5. The locus for Cys373 in Na_v_1.5 is occupied by Tyr or Phe (Tyr362 in Na_v_1.7). (**C**) Critical role of a conserved Tyr in TTX coordination. Shown here is a stereo view of the relative position of Tyr362 and TTX in Na_v_1.7. The aromatic ring of Tyr or Phe at this locus may provide a hydrophobic environment for C4 of TTX. In addition, the bulky ring of Tyr or Phe may stabilize the two Glu residues on P2 helices in repeats I and II, both engaging in TTX binding.

With the reference structure of TTX-bound Na_v_1.5, it becomes clear that one additional role of Tyr362 is to stabilize two invariant Glu residues, Glu364 on repeat I and Glu930 on repeat II in Na_v_1.7, both directly engaging in TTX binding. When the bulky Tyr is replaced by the short side chains of Cys or Ser, there is a large void to confer flexibility for the two Glu residues, which may result in reduced affinity for TTX (Figure 3C). In addition, Tyr362 may also contribute to TTX binding by van der Waals contact with the C4 carbon (Figure 3C). These analyses are compatible with Phe at this locus.

### Structural variations between human Na_v_ channels

Although we have been attempting to capture Na_v_ channels in distinct conformations, it is unsurprising that all the structures of human Na_v_ channels exhibit similar, potentially inactivated conformations. These, disappointing at first glance, actually afford a good opportunity to establish the structure-function relationship of different Na_v_ channels.

When the sequences of the nine human Na_v_ channels are mapped to the structure of Na_v_1.5 using Consurf (32), it is evident that the extracellular segments represent the least conserved region (Figure 4A). Consistently, when the structures of human Na_v_1.2/1.4/1.5/1.7 are superimposed, the extracellular loops deviate the most (Figure 4B). Compared to the rigorously characterized transmembrane region of the pore domain and VSDs, which are responsible for the fundamental functions of any voltage-gated ion channel, the extracellular segments have been underexplored. Comparison of the four human Na_v_ structures in complex with distinct modulators reveals functional significance of the extracellular loops.

**Figure 4.**
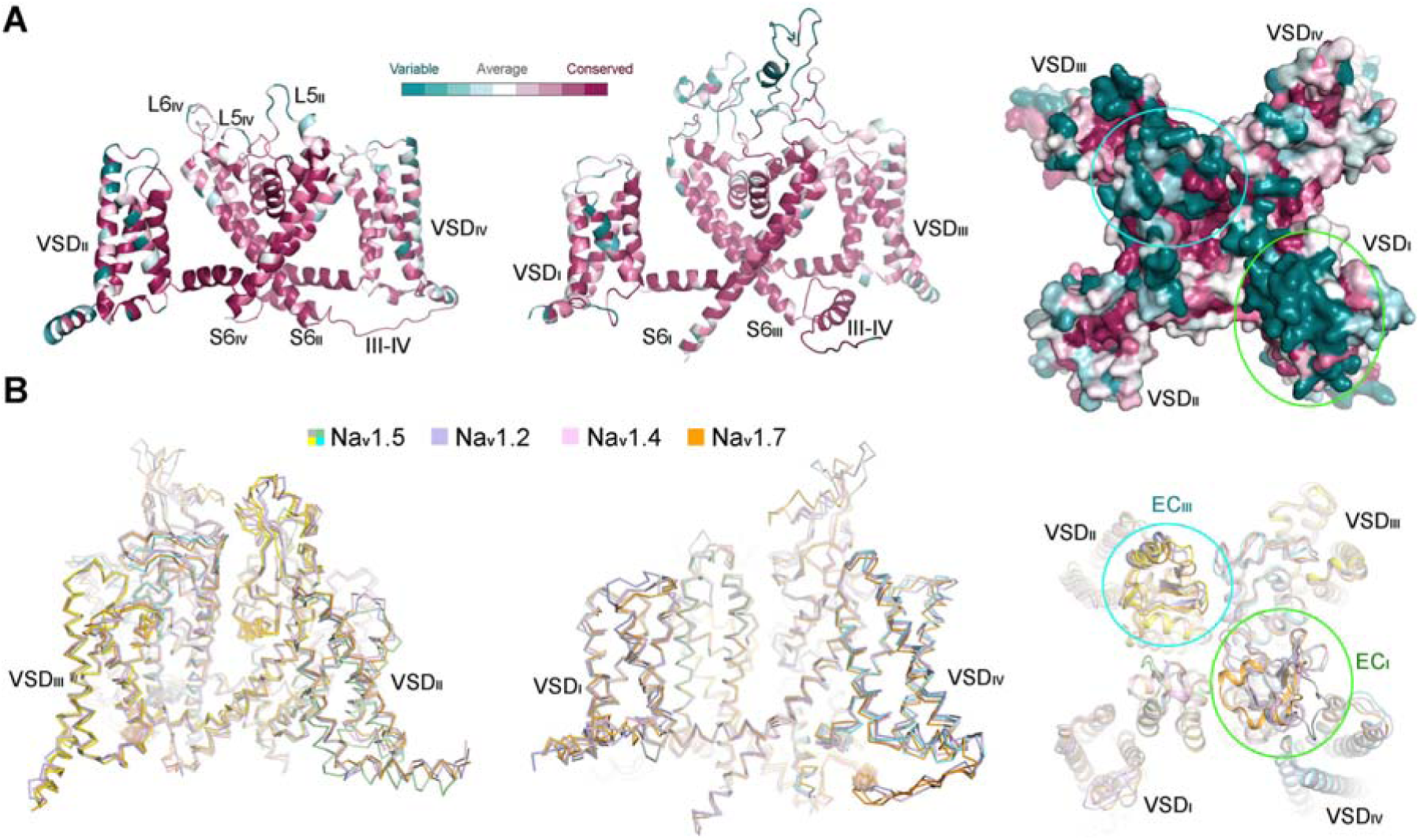
Sequence and structural variations of human Na_v_ channels mainly occur to the extracellular segments. (**A**) Mapping of the conserved residues of human Na_v_ channels to the structure of Na_v_1.5. The extracellular segments represent the most variable regions. Structural mapping of sequence conservation was made in ConSurf (32). *Left and middle*: Side views of the two diagonal repeats. *Right*: Extracellular view of surface presentation of sequence conservation. (**B**) Structures of human Na_v_ channels mainly deviate at the extracellular segments. Structures of human Na_v_1.2/1.4/1.5/1.7, shown as backbone ribbon, are superimposed. The PDB codes are 6J8E for Na_v_1.2, 6AGF for Na_v_1.4, and 6J8J for Na_v_1.7. The blue and green circles in the two panels indicate the corresponding regions that are highly variable.

### Extracellular segments confer specific Na_v_ modulation

The deviation of the extracellular segments is not only in the conformation. Even the secondary structures vary because of the marked difference in length and amino acid composition (Figure 5A). In addition, these segments are enriched of residues for glycosylation and disulfide-bond formation, which add another tier of complexity in the modulation of Na_v_ and the closely related Ca_v_ channels.

**Figure 5.**
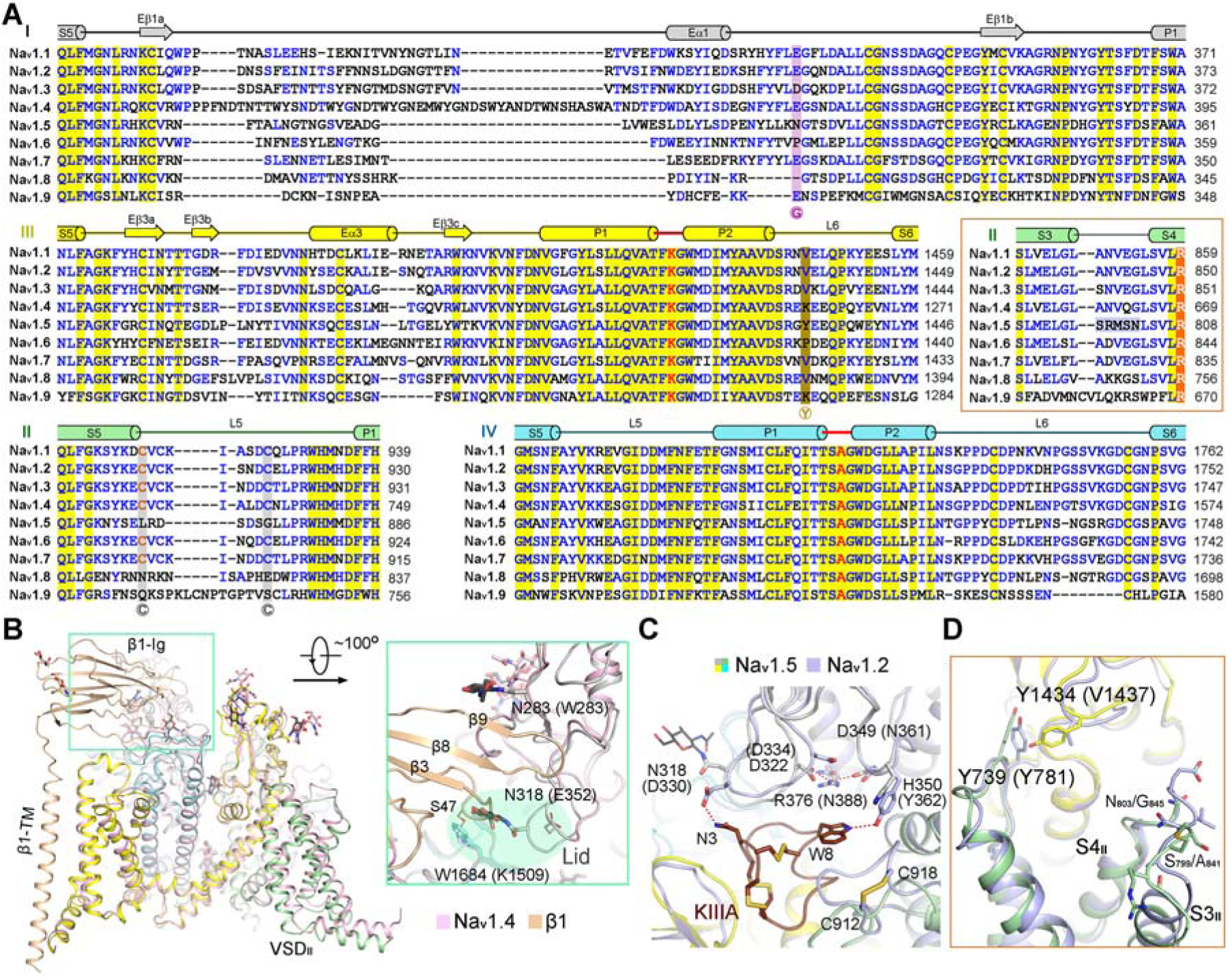
Structural difference of the extracellular segments determines subtype-specific modulation of Na_v_ channels. (**A**) Sequence alignment of the extracellular loops. The boxed comparison is for site 4, the segment between S3 and S4 in VSD_II_. The alignment was adapted from our previous publication with modifications (17). Invariant amino acids are shaded yellow and conserved residues are colored blue. The unique residues that are discussed in panels *B-D* are shaded with dark colors. G: Na_v_1.5-unique glycosylation site (Asn318). Y: Only Na_v_1.5 has a bulky residue (Tyr1434) at this locus. C: The disulfide bond that is conserved in TTX-sensitive Na_v_ channels only. The Uniprot IDs for the aligned sequences are: Na_v_1.1: P35498; Na_v_1.2: Q99250; Na_v_1.3: Q9NY46; Na_v_1.4: P35499; Na_v_1.5: Q14524; Na_v_1.6: Q9UQD0; Na_v_1.7: Q15858; Na_v_1.8: Q9Y5Y9; Na_v_1.9: Q9UI33. (**B**) Structural basis for lack of β1 association with Na_v_1.5. Several factors, particularly the distinct conformation of the “Lid” loop, may prevent Na_v_1.5 from binding to the immunoglobulin (Ig) domain of the β1 subunit. Structures of Na_v_1.5 and β1-bound Na_v_1.4 are superimposed. An enlarged view of the interface between β1-Ig (gold) and the extracellular loops of Na_v_1.4 (pink) and Na_v_1.5 (domain colored) are shown. The corresponding residues in Na_v_1.4 and Na_v_1.5 are shown as thin and thick sticks, and the Na_v_1.4 residues are labeled in brackets. “Lid” refers to the loop that is shaded by semi-transparent green oval. (**C**) Structural basis of the distinct sensitivity of Na_v_1.2 and Na_v_1.5 to μ-conotoxin KIIIA. The corresponding residues in Na_v_1.2 are labeled in brackets. (**D**) Sequence variation in the L6_III_ loop may result in the structural shift of VSD_II_ in Na_v_1.5. Structures of Na_v_1.2 (light purple) and Na_v_1.5 (domain colored) are superimposed and only the described regions are shown in panels C &D.

As described earlier, despite co-expression with the β1 subunit, no density is found for β1 (Figure 1C, Figure S2C). Structural comparison of Na_v_1.5 with the Na_v_1.4-β1 complex reveals the molecular basis for the lack of β1 association (Figure 5B). The primary element is the conformation of the loop following the extracellular helix Eα1, which we will refer to as the Lid loop for its position right above the SF (Figure 5A,B). The Lid is positioned lower toward the SF in Na_v_1.5 than in Na_v_1.4, resulting in a potential steric hindrance with the hairpin loop between the β2 and β3 strands in the β1 subunit (Figure 5B, inset). More importantly, the residue at the tip of the lid loop is Asn318, a glycosylation site unique to Na_v_1.5 among the nine subtypes (Figure 5A,B). The sugar moieties would clash with β1. Upward movement of the loop, which may potentially change the position of Asn318 and the attached glycan to avoid the steric clash, is restricted by the interaction between Arg376 on P2_I_ and Asp322 on the Lid. Note that the locus for Arg376 is occupied by Arg or Lys in TTXr and Asn in TTXs channels (Figure 5A). There is no interaction between the corresponding Asn and Lid residues in Na_v_1.2/1.4/1.7, hence Lid can be liberated to adopt a higher position (Figure 5C).

The extracellular region harbors the primary binding sites for various peptidic toxins in addition to the β subunits (18, 20). Conformational change of the extracellular loops thereby also underscores the varied toxin sensitivity by different Na_v_ subtypes. For instance, it has been shown that Na_v_1.2 can be blocked by μ-conotoxin KIIIA almost irreversibly, whereas the IC_50_ for the rat Na_v_1.5 is ∼ 284 μM (33). Structure-based sequence analysis has revealed three KIIIA binding residues in Na_v_1.2 that are altered in Na_v_1.5, Asp330→Asn318, Tyr362→His350, and Tyr1443→Trp1440 (18). Structural comparison between Na_v_1.5 and KIIIA-bound Na_v_1.2 affords additional insight for the distinct sensitivity toward KIIIA.

The Tyr1443→Trp1440 variation may have little or no effect on KIIIA binding, because the π-cation interaction is preserved. Tyr362→His350 can lead to reduced affinity because of the loss of one hydrogen bond with Trp8 on KIIIA (Figure 5C). Glycosylated Asn318 by the residue *per se* can still form a hydrogen bond with KIIIA-Asn3; however, it is the conformation of the lid loop that leaves it out of reach of interacting with any residue on KIIIA (Figure 5C). Last but not least, the L5_II_ loop, a short loop between S5 and P1 helix in repeat II, exhibits different conformations because Na_v_1.5 lacks a disulfide bond that is only conserved in TTXs subtypes (Figure 5A,C). The conformational shift of L5_II_ may also contribute to the affinity difference with KIIIA.

When the four human Na_v_ structures are superimposed, VSD_II_ of Na_v_1.5 deviates from the other three by slight rotation, although it also exhibits a similar “up” conformation as VSD_II_s in the other three channels (Figures 4B,5B). The sequences of the transmembrane segments are highly conserved between Na_v_1.5 and the other three Na_v_ channels (17). The structural shifts are thus likely owing to the interaction between VSD_II_ and other segments. As the transmembrane helices are nearly unchanged, we examined the interface at the extracellular region. Na_v_1.5 has a unique Tyr1434 on L6_III_, a locus that is occupied by Val in Na_v_1.2/1.5 and Lys in Na_v_1.4 (Figure 5A). Because of this bulky residue, a conserved Tyr on the S1-S2 loop in VSD_II_ has to shift slightly. Although it is only a relatively minor change, this Tyr residue (Tyr739 in Na_v_1.5) appears to be the pivoting center for the overall shift of VSD_II_. Therefore, the positional change of Na_v_1.5-VSD_II_ is in part because of an allosteric result from the alternation of the extracellular segments (Figure 5D).

We cannot exclude the contribution of other factors that are invisible in the present 3D reconstruction. For instance, site 4, i.e., the extracellular segments of S3 and S4 and the intervening linker of VSD_II_, is completely different between Na_v_1.5 and the other three Na_v_ channels (Figure 5A, inset). Accordingly, the structure of the S3-S4 loop in Na_v_1.5 deviates from that in Na_v_1.2. Its interaction with the indiscernible detergents or lipids may also contribute to the positional shift of VSD_II_. In addition, the invisible intracellular segments may also play a role for the conformational deviation of VSD_II_.

## Discussion

In this paper, we report the structures of human Na_v_1.5 bound to TTX or quinidine. As this is the fourth human Na_v_ channel whose structure has been elucidated, we hereby try to avoid describing the common structural features and mechanisms that have been previously described in detail, including the overall architecture, Na^+^ path through SF, VSDs, fast inactivation mechanism, intracellular gating and fenestrations. Instead, we focus on comparative structural analysis in the hope to provide an advanced understanding of the uniqueness of different Na_v_ channels.

During this manuscript preparation, cryo-EM structures of a rat Na_v_1.5 variant that contains truncations in the I-II linker, the II-III linker, and the C-terminus was reported in complex with TTX and flecainide, a class Ic antiarrhythmic drug (34). There are only two overlapping points between the published paper and our present study, the molecular basis for TTX-resistance and the lack of β1 association. Of particular note, we have different interpretations for the same structural observations.

Jiang and co-workers attributed the role of Tyr for TTX binding in TTXs channels as π-π packing, which has not been observed in our reported structures of TTX-bound Na_v_PaS or Na_v_1.7. In Figure 3B, we provide a detailed analysis. With respect to β1 association, both studies identified the glycosylated Asn318 to impede β1 binding. However, our analysis indicates that the conformation of the Lid loop, which is locked by Arg376 in Na_v_1.5, determines the specific position of Asn318. If the Lid loop in Na_v_1.5 can exhibit the same conformation as that in Na_v_1.2/1.4/1.7, Asn318 would be oriented to an opposite direction that has no collision with β1, as seen for Glu352 in Na_v_1.4 (Figure 5B, inset).

In addition to the differences in interpreting the same structural observation, we hereby mainly focus on the structural distinctions between Na_v_1.5 and the previously reported Na_v_ structures. Comparative analysis has identified the extracellular loops above the pore domain to be important determinants for subtype-specific channel modulation by different auxiliary subunits and toxins, and potentially other channel properties. Supporting the functional significance of the extracellular loops, nearly 40 mutations associated with LQT3 or Brugada syndrome are mapped to the extracellular loops (Tables S1, S2). Most of these residues are well-resolved. Although some of the mutations, such as C1363Y and C1728R/W, may compromise structural stability because of the disruption of disulfide bonds, the functional relevance of the majority of these disease residues is not immediately clear, as they do not participate in intra-protein interactions. It remains to be investigated whether the pathogenic mechanism of these residues involves binding factors or channel folding and membrane trafficking.

In sum, elucidation the cryo-EM structures of human Na_v_ channels from different tissues affords unprecedented opportunity for establishment of the structure-function relationship.

## Acknowledgements

We thank Xiaomin Li (Tsinghua University) for technical support during EM image acquisition. This work was funded by the National Key R&D Program (2016YFA0500402 to X.P.) from Ministry of Science and Technology of China, and the National Natural Science Foundation of China (projects 31621092, 31630017, and 81861138009). We thank the Tsinghua University Branch of China National Center for Protein Sciences (Beijing) for providing the cryo-EM facility support. We thank the computational facility support on the cluster of Bio-Computing Platform (Tsinghua University Branch of China National Center for Protein Sciences Beijing) and the “Explorer 100” cluster system of Tsinghua National Laboratory for Information Science and Technology. N.Y. is supported by the Shirley M. Tilghman endowed professorship from Princeton University.

## Author contributions

N.Y. conceived the project. Z.L., G.H., X.P., and J.L. performed experiments for structural determination. X.J., K.W., and X.P. performed and analyzed electrophysiological measurements. All authors contributed to data analysis. N.Y. wrote the manuscript.

