## Supplementary material for "Structural basis for pore blockade of the human cardiac sodium channel Na_v_1.5 by tetrodotoxin and quinidine": Materials and methods; supplementary Figures 1-4, Tables 1-5

#### Transient co-expression of human $Na_v1.5$ and β1

The optimized coding DNAs for human $Na_v1.5$ (Uniprot: Q14524) was cloned into the pEG BacMam vector with twin Strep-tag and FLAG tag in tandem at the amino terminus, and β1 (Uniprot: Q07699) was cloned into the pCAG vector without affinity tag (1, 2). HEK293F cells (Invitrogen) were cultured in SMM 293T-I medium (Sino Biological Inc.) under 5% $CO_2$ in a Multitron-Pro shaker (Infors, 130 r.p.m.) at 37 $^oC$, and transfected with plasmids when the cell density reached $2 \times 10^6$ cells per ml. For one liter cell culture, approximately 2 mg plasmids (1.4 mg for $Na_v1.5$ plus 0.6 mg for β1) were pre-incubated with 4 mg 25-kDa linear polyethylenimines (PEIs) (Polysciences) in 30 mL fresh medium for 15-30 min before adding into cell culture. Transfected cells were cultured for 48 h before harvesting.

#### Purification of $Na_v1.5$ and β1 protein

16 L transfected cells were harvested by centrifugation at 800 g and resuspended in the lysis buffer containing 25 mM Tris-HCl (pH 7.5) and 150 mM NaCl. The suspension was supplemented with 1% (w/v) n-dodecyl-β-D-maltopyranoside (DDM, Anatrace), 0.1% (w/v) cholesteryl hemisuccinate Tris salt (CHS, Anatrace), and protease inhibitor cocktail containing 2 mM phenylmethylsulfonyl fluoride (PMSF), aprotinin (3.9 μg/ml), pepstatin (2.1 μg/ml), and leupeptin (15 μg/ml). After incubation at 4 $^oC$ for 3 h, the cell lysate was ultra-centrifuged at 20,000 g for 45 min, and the supernatant was applied to anti-Flag M2 affinity gel (Sigma) by gravity at 4 $^oC$. The resin was rinsed four times with the W buffer that contains 25 mM Tris-HCl (pH 7.5), 150 mM NaCl, 0.06% glycol-diosgenin (GDN, Anatrace), and the protease inhibitor cocktail. Proteins were eluted with W buffer plus 200 μg/ml FLAG peptide (Sigma). The eluent was then applied to Strep-Tactin Sepharose (IBA) and the purification protocol was similar to the previous steps except elution buffer, which was W buffer plus 2.5 mM D-Desthiobiotin (IBA). The eluent was then concentrated using a 100-kDa cut-off Centricon (Millipore) and further purified by Superose-6 column (GE Healthcare) in W buffer. The presence of target proteins was confirmed

by mass spectrometry. The purified proteins were pooled and concentrated to approximately 2 mg/ml for cryo-EM analysis. TTX (66 μM) and untagged calmodulin (CaM, ~200 μM) or quinidine (Target Mol, 1 mM) and STX (12 μM) were separately added to the concentrated sample one hour before cryo grids preparation.

**Whole cell electrophysiology**

HEK293T cells cultured in Dulbecco's Modified Eagle Medium (DMEM, BI) supplemented with 4.5 mg/ml glucose and 10% fetal bovine serum (FBS, BI) were plated onto glass coverslips for subsequent patch clamp recordings. Cells were transiently co-transfected using lipofectamine 2000 (Invitrogen) with the expression plasmids for indicated $Na_v1.5$ WT (with or without β1) or mutants and an eGFP-encoding plasmid. Cells with green fluorescence were selected for patch-clamp recording at 18–36 h after transfection. All experiments were performed at room temperature. No further authentication was performed for the commercially available cell line. Mycoplasma contamination was not tested.

The whole-cell $Na^+$ currents were recorded in HEK293T cells using an EPC10-USB amplifier with Patchmaster software v2*90.2 (HEKA Elektronic), filtered at 3 kHz (low-pass Bessel filter) and sampled at 50 kHz. The borosilicate pipettes (Sutter Instrument) used had a resistance of 2-4 MΩ and the electrodes were filled with the internal solution composed of (in mM) 105 CsF, 40 CsCl, 10 NaCl, 10 EGTA, 10 HEPES, pH 7.4 with CsOH. The bath solutions contained (mM): 140 NaCl, 4 KCl, 10 HEPES, 10 D-Glucose, 1 MgCl2, 1.5 CaCl2, pH 7.4 with NaOH. Data were analyzed using Origin (OriginLab) and GraphPad Prism (GraphPad Software).

The voltage dependence of ion current (I-V) was analyzed using a protocol consisting of steps from a holding potential of -120 mV (for 100 ms) to voltages ranging from -90 to +80 mV for 50 ms in 5 mV increment. The linear component of leaky currents and capacitive transients were subtracted using the P/4 procedure. In the activation and conductance density calculation, we used the equation, $G = I/(V-V_r)$, where Vr (the reversal potential) represents the voltage at which the current is zero. For the activation curves, conductance (G) was normalized and plotted against the voltage from -90 mV to +20 mV. To obtain the conductance density curves, G was divided by the capacitance (C), and plotted against the voltage from -90 mV to +20 mV. For voltage dependence of inactivation, cells were clamped at a holding potential of -90 mV, and were applied to step pre-pulses from -120 mV to +20 mV for 100 ms with an increment of 5 mV. Then, the $Na^+$ current was recorded at the test pulse of 0 mV for 50 ms. The peak currents under the test pulses were normalized and plotted against the pre-pulse voltage. Activation and inactivation curves were fit to a Boltzmann function to obtain $V_{1/2}$ and slope values. Time course of inactivation data from the peak current at 0 mV was fitted to a single exponential equation: $y = A1 \exp(-x/\tau_{inac}) + y0$, where A1 was the relative fraction of current inactivation, $\tau_{inac}$ was the time constant, x was the time, and y0 was the amplitude of the steady-state component.

To investigate use-dependent blockade of $Na_v$1.5 variants by quinidine, currents were recorded after 50-ms pulses at 1 Hz from -120 mV to 0 mV. Solutions with different quinidine concentrations were perfused to the recording cell using a multichannel perfusion system (VM8, ALA). Concentration-response curve was fitted with: Y = Bottom + (Top - Bottom) / (1 + 10^((Log$IC_{50}$-X) * HillSlope)), where $IC_{50}$ is the concentration of quinidine that blocks 50% of the current and X is log of quinidine concentration, and HillSlope is slope factor. All data points are presented as mean ± standard error of the mean (SEM) and n is the number of experimental cells from which recordings were obtained. Statistical significance was assessed using an unpaired t-test with Welch's correction and extra sum-of-squares F test.

**Cryo-EM data acquisition**

Aliquots of 3.5 μl freshly purified $Na_v$1.5-β1 were placed on glow-discharged holey carbon grids (Quantifoil Au 300 mesh, R1.2/1.3). Grids were blotted for 3.0 s and plunge frozen in liquid ethane cooled by liquid nitrogen with Vitrobot Mark IV (Thermo Fisher). Electron micrographs were acquired on a Titan Krios electron microscope (Thermo Fisher) operating at 300 kV and equipped with Cs corrector, Gatan K2 Summit detector and GIF Quantum energy filter. A total of 5,780 and 7,799 movie stacks were automatically collected for $Na_v$1.5T (with TTX) and $Na_v$1.5Q (with quinidine), respectively, using AutoEMation (3) with a slit width of 20 eV on the energy filter and a preset defocus range from -1.8 µm to -1.5 µm in super-resolution mode at a nominal magnification of 105,000X. Each stack was exposed for 5.6 s with 0.175 s per frame, resulting in 32 frames per stack. The total dose rate was 48 $e^{-}/Å^{2}$ for each stack. The stacks were motion corrected with MotionCor2 (4) and binned 2-fold, resulting in 1.091 Å/pixel. Meanwhile, dose weighting was performed (5). The defocus values were estimated with Gctf (6).

**Image processing**

A diagram for the workflow of data processing is presented in Figure S2. A total of 3,011,243 and 3,979,224 particles were automatically picked using RELION (7-9) from 5,780 and 7,799 collected micrographs for $Na_v$1.5T and $Na_v$1.5Q, respectively. After 2D classification, 708,605 good particles for $Na_v$1.5T and 1,186,235 good particles for $Na_v$1.5Q were selected and applied to global angular searching 3D classification with K set to 1 (one class). For each of the last several iterations of the global angular searching 3D classification, a local angular searching 3D classification was performed with 4 classes. A total of non-redundant 231,131 particles for $Na_v$1.5T and 320,122 particles for $Na_v$1.5Q were selected from the local angular searching 3D classification and subjected to 3 cycles of multi-reference 3D classification to remove bad particles. Then 145,139 and 124,954 selected particles were subject to 3D auto-refinement, yielding 3D maps with overall resolutions of 3.4 Å for $Na_v$1.5T and 3.3 Å for $Na_v$1.5Q, which were further improved to 3.3 Å for $Na_v$1.5T and 3.2 Å for $Na_v$1.5Q after application of a soft mask. 2D classification, 3D classification, and auto-refinement were performed in RELION 3.0. Resolution was estimated with the gold-standard Fourier shell correlation 0.143 criterion (10) with high resolution noise substitution (11).

**Model building and structure refinement**

Sequences for the nine human $Na_v$ channels were aligned using Clustal W. Model building was performed based on the 3.4 Å reconstruction map for $Na_v$1.5T. The coordinates of human $Na_v1.4$ (PDB accession number: 6AGF) were fitted into the EM map by CHIMERA(12). The sequence of human $Na_v1.4$ were mutated to corresponding residues in human $Na_v1.5$ in COOT (13) and each residue was manually checked. The chemical properties of amino acids were considered during model building. There was no density corresponding to β1 in either map. STX was not modelled in $Na_v1.5Q$ due to low resolution.

In both structures, 1,151 side chains and 9 sugar moieties were built for human $Na_v1.5$. The N-terminal 118 residues, intracellular I-II linker (residues 430-698), II-III linker (residues 945-1187), and C-terminal sequences after Ser1782 cannot be resolved. TTX and one GDN molecule were modelled in $Na_v1.5T$, and quinidine was assigned in $Na_v1.5Q$.

Structure refinement was performed using phenix.real_space_refine application in PHENIX (14) real space with secondary structure and geometry restraints. Over-fitting of the overall model was monitored by refining the model in one of the two independent maps from the gold-standard refinement approach and testing the refined model against the other map (15). Statistics of the map reconstruction and model refinement can be found in Table S5.

### Supplementary Figures and Legends

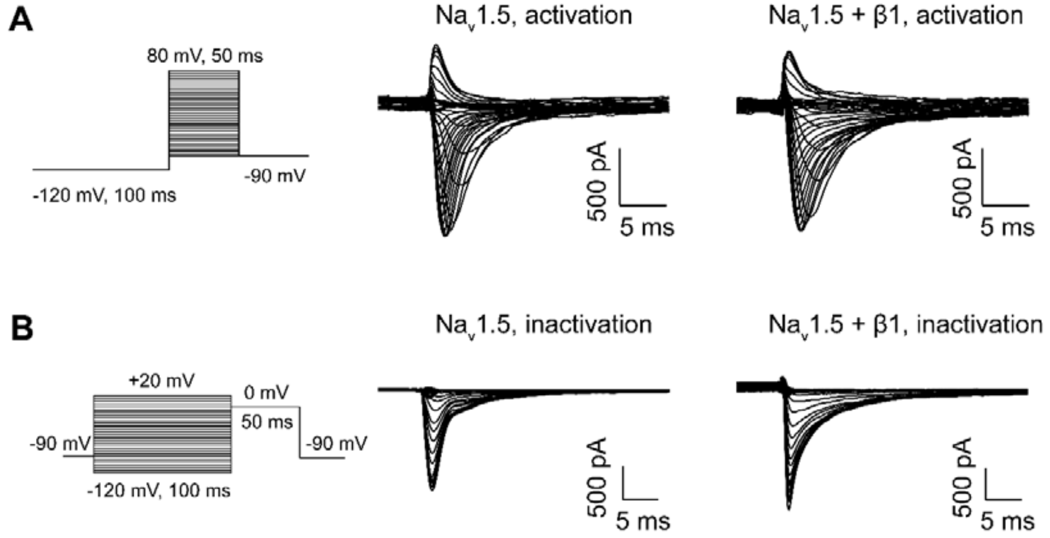


**Supplementary Figure 1 | Electrophysiological properties of Na$_v$1.5 transiently co-expressed with or without β1 subunits in HEK293T cells.**

(**A**) Voltage-dependent activation traces. (**B**) Voltage-dependent inactivation traces. The left panels show the diagrams of recording protocols. The traces for wild type Na$_v$1.5 and Na$_v$1.5 + β1 in the presence of 100 nM TTX are shown in the middle and right panels, respectively. Please refer to Methods for details.

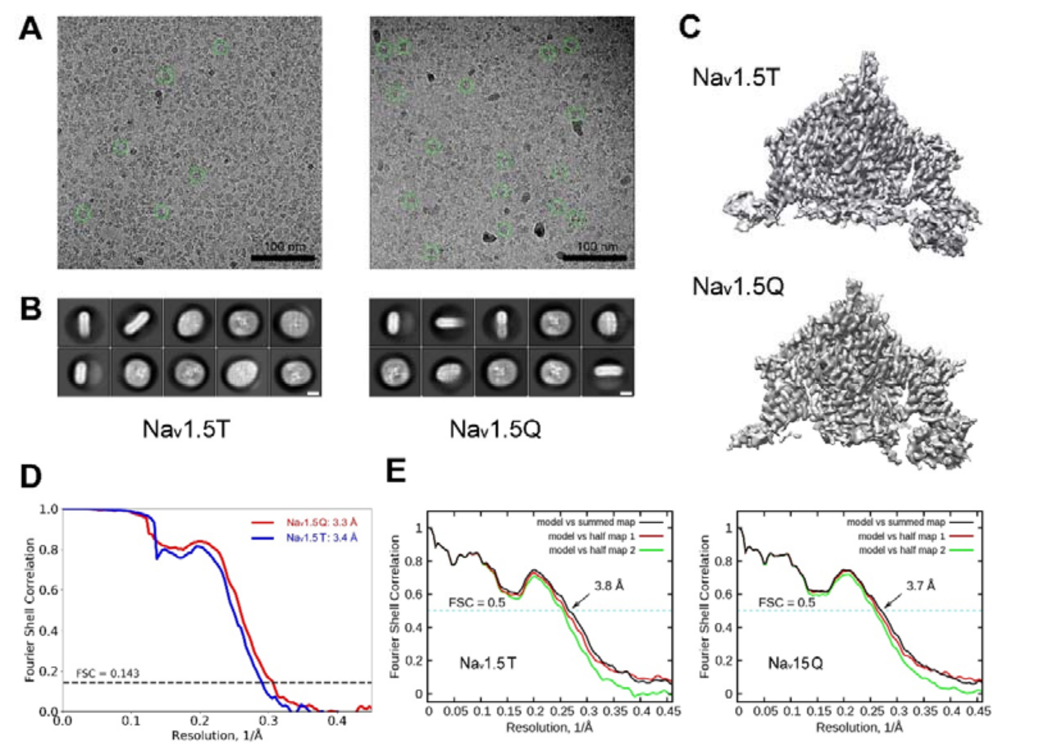


**Supplementary Figure 2 | Cryo-EM analysis of human $Na_v$1.5.**

(**A**) Representative electron micrographs of $Na_v$1.5 bound to TTX ($Na_v$1.5T, left) and quinidine ($Na_v$1.5Q, right). Particles in distinct orientations are highlighted by green circles. Scale bar represents 100 nm. (**B**) Representative two-dimensional class averages of $Na_v$1.5T and $Na_v$1.5Q. White scale bars represent 5 nm. (**C**) The 3D EM reconstructions of $Na_v$1.5T and $Na_v$1.5Q. The maps were generated in CHIMERA (12). (**D**) Gold-standard Fourier shell correlation (FSC) curve for the 3D reconstruction of $Na_v$1.5T and $Na_v$1.5Q. (**E**) FSC curves of the refined model versus the overall map that is was refined against (black), of the model refined in the first of the two independent maps used for the gold-standard FSC versus that same map (red), and of the model refined in the first of the two independent maps versus the second independent map (green). The small difference between the red and green curves indicates that the refinement of the atomic coordinates did not suffer from overfitting.

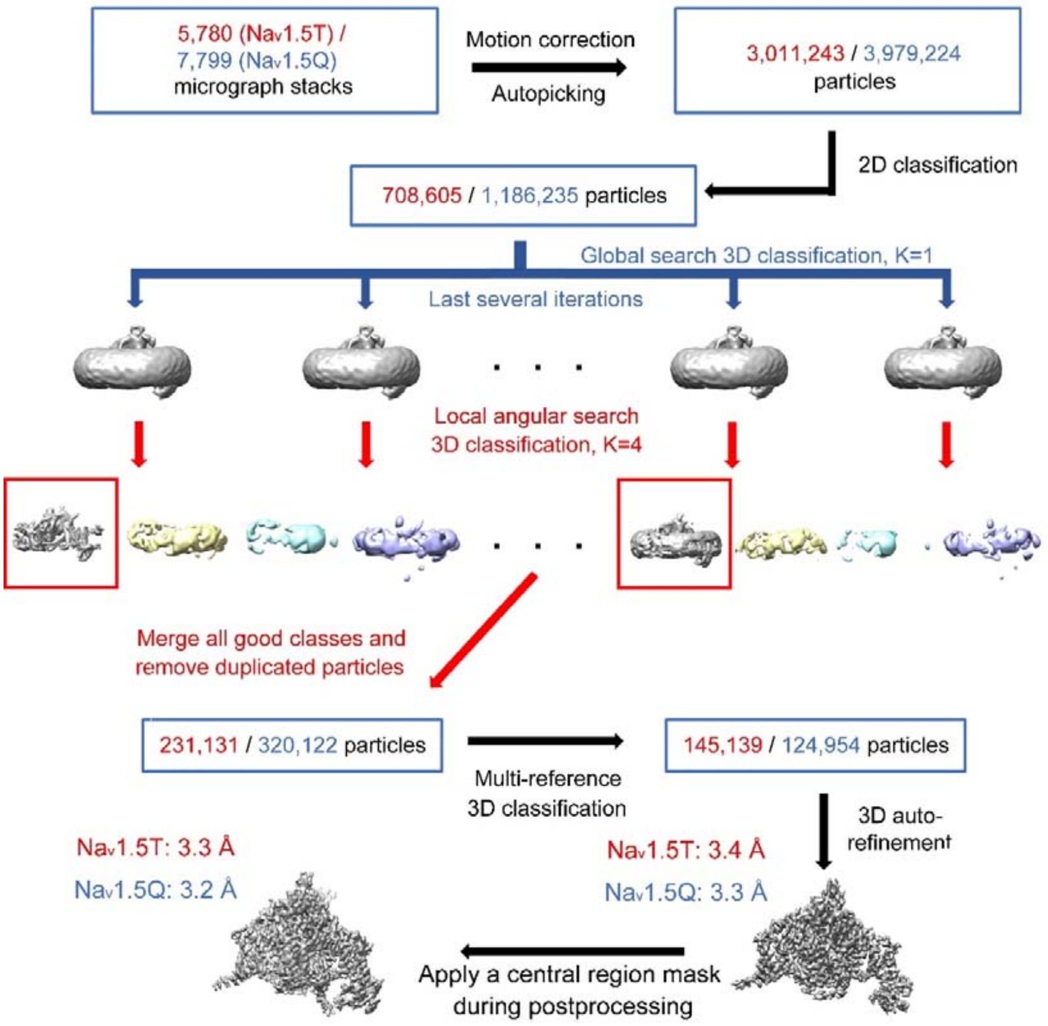


**Supplementary Figure 3 | The flowchart for EM data processing.**

Details can be found in Materials and Methods.

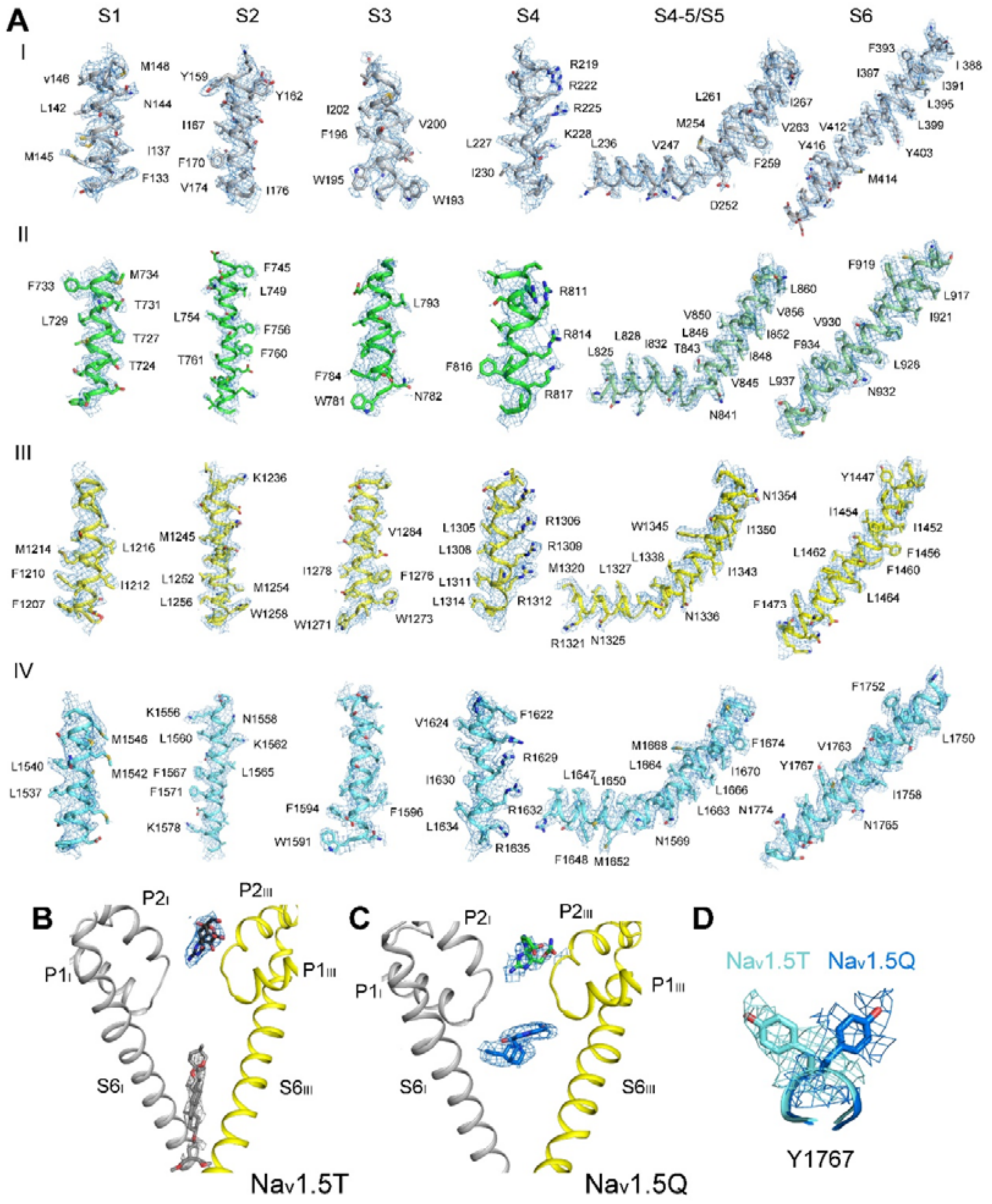


**Supplementary Figure 4 | EM maps for representative segments of Na$_v$1.5T and Na$_v$1.5Q.**

(**A**) EM maps for the S1-S6 segments in each repeat of Na$_v$1.5T. (**B**) EM maps for TTX and GDN of Na$_v$1.5T. (**C**) EM maps for STX and quinidine of Na$_v$1.5Q. (**D**) Conformational change of Y1767 in Na$_v$15T and Na$_v$1.5Q. The maps were prepared in PyMol (16) and contoured at 3-4 σ.

**Supplementary Table 1 | Mutations on human $Na_v$1.5 that are associated with long-QT syndrome, type 3.**

| Mutations | Structure |
|---|---|
| G9V | NTD |
| R18Q | NTD |
| R27H | NTD |
| E30G | NTD |
| R43Q | NTD |
| E48K | NTD |
| P52S | NTD |
| R53Q | NTD |
| R104G | NTD |
| S115G | NTD |
| V125L | NTD |
| L212P | S3-4I |
| R222Q | S4I |
| R225Q | S4I |
| R225W | S4I |
| V240M | S4-5I |
| Q245K | S4-5I |
| V247L | S4-5I |
| N275K | L5I |
| G289S | L5I |
| R340W | L5I |
| R367C | SF I |
| T370M | SF I |
| I397T | S6I |
| L404Q | S6I |
| N406K | S6I |
| L409V | S6I |
| V411M | S6I |
| A413E | S6I |
| A413T | S6I |
| E462A | I-II linker |
| E462K | I-II linker |
| F530V | I-II linker |
| R535Q | I-II linker |
| R569W | I-II linker |
| S571I | I-II linker |
| A572D | I-II linker |
| A572S | I-II linker |
| A572V | I-II linker |
| Q573E | I-II linker |
| G579R | I-II linker |
| G615E | I-II linker |
| L619F | I-II linker |
| P637L | I-II linker |
| G639R | I-II linker |
| P648L | I-II linker |
| E654K | I-II linker |
| L673P | I-II linker |
| R680H | I-II linker |
| R689C | I-II linker |
| R689H | I-II linker |
| P701L | I-II linker |
| T731I | S1II |
| Q750R | S2II |
| D772N | S2-3II |
| F816Y | S4II |
| I848F | S5II |
| S941N | S6II |
| Q960K | II-III linker |
| R965L | II-III linker |
| R971C | II-III linker |
| C981F | II-III linker |
| A997S | II-III linker |
| C1004R | II-III linker |
| E1053K | II-III linker |
| T1069M | II-III linker |
| A1100V | II-III linker |
| D1114N | II-III linker |
| D1166N | II-III linker |
| R1193Q | II-III linker |
| Y1199S | II-III linker |
| E1225K | S1-2III |
| E1231K | S1-2III |
| F1250L | S2III |
| L1283M | S3III |
| E1295K | S3-4III |
| T1304M | S4III |
| N1325S | S4-5III |
| A1326S | S4-5III |
| A1330P | S4-5III |
| A1330T | S4-5III |
| P1332L | S5III |
| S1333Y | S5III |
| I1334V | S5III |
| L1338V | S5III |
| R1432S | L6III |
| S1458Y | S6III |
| N1472S | S6III |
| F1473C | S6III |
| G1481E | III-IV linker |
| F1486L | III-IV linker |
| M1487L | III-IV linker |
| T1488R | III-IV linker |
| E1489D | III-IV linker |
| K1493R | III-IV linker |
| Y1495S | III-IV linker |
| M1498V | III-IV linker |
| L1501V | III-IV linker |
| K1505N | III-IV linker |
| V1532I | S1IV |
| L1560F | S2IV |
| I1593M | S3IV |
| F1594S | S3IV |
| F1596I | S3IV |
| S1609W | S3IV |
| T1620K | S4IV |
| R1623L | S4IV |
| R1623Q | S4IV |
| R1626H | S4IV |
| R1626P | S4IV |
| R1644C | S4-5IV |
| R1644H | S4-5IV |
| T1645M | S4-5IV |
| L1650F | S4-5IV |

| Mutations | Structure |
|---|---|
| M1652R | S4-5IV |
| M1652T | S4-5IV |
| I1660V | S5IV |
| V1667I | S5IV |
| T1723N | L6IV |
| R1739W | L6IV |
| L1761F | S6IV |
| L1761H | S6IV |
| V1763M | S6IV |
| M1766L | S6IV |
| Y1767C | S6IV |
| I1768V | S6IV |
| V1777M | S6IV |
| T1779M | S6IV |

| Mutations | Structure |
|---|---|
| E1784K | CTD |
| D1790G | CTD |
| Y1795C | CTD |
| Y1795YD | CTD |
| D1819N | CTD |
| L1825P | CTD |
| R1826H | CTD |
| D1839G | CTD |
| H1849R | CTD |
| R1897W | CTD |
| E1901Q | CTD |
| S1904L | CTD |
| Q1909R | CTD |

| Mutations | Structure |
|---|---|
| R1913H | CTD |
| A1949S | CTD |
| V1951L | CTD |
| Y1977N | CTD |
| F2004L | CTD |
| F2004V | CTD |
| R2012C | CTD |
| ΔA586-L587 | I-II linker |
| ΔE429 | I-II linker |
| ΔI1212 | S1III |
| ΔK1505-Q1507 | III-IV linker |
| ΔQ1507-P1509 | III-IV linker |
| ΔF1617 | S3-4IV |

Disease mutations that are mapped to the extracellular segments are shaded with their respective domain colors, and structurally unresolved regions are shaded light gray. The same applies to Tables S2 and S3.

Mutations in Tables S1-S3 are summarized from https://www.uniprot.org/uniprot/Q14524

**Supplementary Table 2 | Mutations on human $Na_v1.5$ that are associated with Brugada syndrome.**

| Mutations | Structure | Mutations | Structure | Mutations | Structure |
|---|---|---|---|---|---|
| R18Q | NTD | L315P | L5I | P701L | I-II linker |
| R27H | NTD | G319S | L5I | P717L | I-II linker |
| N70K | NTD | T320N | L5I | A735E | S1II |
| D84N | NTD | L325R | L5I | A735V | S1II |
| F93S | NTD | P336L | L5I | E746K | S1-2II |
| I94S | NTD | G351D | L5I | G752R | S2II |
| V95I | NTD | G351V | L5I | G758E | S2II |
| R104Q | NTD | T353I | L5I | M764R | S2II |
| R104W | NTD | D356N | L5I | D772N | S2-3II |
| N109K | NTD | R367C | SF I | P773S | S2-3II |
| R121Q | NTD | R367H | SF I | V789I | S3II |
| R121W | NTD | R367L | SF I | R808P | S4II |
| K126E | NTD | M369K | SF I | L812Q | S4II |
| L136P | S1I | W374G | SF I | R814Q | S4II |
| V146M | S1I | R376H | SF I | K817E | S4II |
| E161K | S2I | G386E | L6I | L839P | S5II |
| E161Q | S2I | G386R | L6I | F851L | S5II |
| K175N | S2I | V396A | S6I | E867Q | L5II |
| A178G | S2I | V396L | S6I | R878C | L5II |
| C182R | S2-3I | N406S | S6I | R878H | L5II |
| A185V | S2-3I | E439K | I-II linker | H886P | SF II |
| T187I | S2-3I | D501G | I-II linker | F892I | SF II |
| A204V | S3I | G514C | I-II linker | R893C | SF II |
| L212Q | S3-4I | R526H | I-II linker | R893H | SF II |
| T220I | S4I | F532C | I-II linker | C896S | SF II |
| R222Q | S4I | F543L | I-II linker | E901K | SF II |
| V223L | S4I | G552R | I-II linker | S910L | L6II |
| A226V | S4I | L567Q | I-II linker | C915R | S6II |
| I230V | S4I | G615E | I-II linker | L917R | S6II |
| V232I | S4-5I | L619F | I-II linker | N927S | S6II |
| V240M | S4-5I | R620C | I-II linker | L928P | S6II |
| Q270K | S5I | T632M | I-II linker | L935P | S6II |
| L276Q | L5I | P640A | I-II linker | R965C | II-III linker |
| H278D | L5I | A647D | I-II linker | R965H | II-III linker |
| R282C | L5I | P648L | I-II linker | A997T | II-III linker |
| R282H | L5I | R661W | I-II linker | R1023H | II-III linker |
| V294M | L5I | H681P | I-II linker | E1053K | II-III linker |
| V300I | L5I | C683G | I-II linker | D1055G | II-III linker |

| Mutations | Structure | Mutations | Structure | Mutations | Structure |
|---|---|---|---|---|---|
| S1079Y | II-III linker | L1412F | SF III | G1661R | S5IV |
| A1113V | II-III linker | K1419E | SF III | V1667I | S5IV |
| S1140T | II-III linker | G1420R | SF III | S1672Y | S5IV |
| R1193Q | II-III linker | A1427S | SF III | A1680T | L5IV |
| S1219N | S1III | A1428V | SF III | D1690N | L5IV |
| E1225K | S1-2III | R1432G | L6III | A1698T | SF IV |
| Y1228H | S1-2III | R1432S | L6III | T1709M | SF IV |
| R1232Q | S1-2III | G1433V | L6III | T1709R | SF IV |
| R1232W | S1-2III | P1438L | L6III | G1712S | SF IV |
| K1236N | S2III | E1441Q | L6III | D1714G | SF IV |
| L1239P | S2III | I1448L | S6III | N1722D | L6IV |
| D1243N | S2III | I1448T | S6III | C1728R | L6IV |
| V1249D | S2III | Y1449C | S6III | C1728W | L6IV |
| E1253G | S2III | V1451D | S6III | G1740R | L6IV |
| G1262S | S2-3III | N1463Y | S6III | G1743E | L6IV |
| W1271C | S3III | V1468F | S6III | G1743R | L6IV |
| A1288G | S3III | Y1494N | III-IV linker | G1748D | S6IV |
| F1293S | S3-4III | L1501V | III-IV linker | V1764F | S6IV |
| L1311P | S4III | G1502S | III-IV linker | T1779M | S6IV |
| G1319V | S4-5III | R1512W | III-IV linker | E1784K | CTD |
| V1323G | S4-5III | I1521K | III-IV linker | Y1795H | CTD |
| P1332L | S4-5III | V1525M | III-IV linker | Y1795YD | CTD |
| F1344L | S5III | K1527R | III-IV linker | Q1832E | CTD |
| F1344S | S5III | E1548K | S1-2IV | C1850S | CTD |
| L1346S | S5III | A1569P | S2IV | V1861I | CTD |
| L1346P | S5III | F1571C | S2IV | K1872N | CTD |
| M1351R | S5III | E1574K | S2IV | V1903L | CTD |
| V1353M | S5III | L1582P | S2-3IV | A1924T | CTD |
| G1358W | L5III | R1583C | S2-3IV | G1935S | CTD |
| K1359N | L5III | R1583H | S2-3IV | E1938K | CTD |
| F1360C | L5III | V1604M | S3IV | V1951L | CTD |
| C1363Y | L5III | Q1613L | S3-4IV | I1968S | CTD |
| S1382I | L5III | T1620M | S3-4IV | F2004L | CTD |
| V1405L | SF III | R1623Q | S4IV | F2004V | CTD |
| V1405M | SF III | R1629Q | S4IV | ΔF393 | S6I |
| G1406E | SF III | G1642E | S4-5IV | ΔK1479 | III-IV linker |
| G1406R | SF III | R1644C | S4-5IV | ΔK1500 | III-IV linker |
| G1408R | SF III | A1649V | S4-5IV | ΔF1617 | S3-4IV |
| Y1409C | SF III | I1660V | S5IV | | |

**Supplementary Table 3 | Mutations on human $Na_v1.5$ that are associated with cardiac disorders other than long QT syndrome and Brugada syndrome.**

| Mutations | Disease | Structure |
|---|---|---|
| E161K | PFHB1A | S2I |
| R225W | PFHB1A | S4I |
| G298S | PFHB1A | L5I |
| T512I | PFHB1A | I-II linker |
| G514C | PFHB1A | I-II linker |
| G752R | PFHB1A | S2II |
| R1232W | PFHB1A | S1-2III |
| D1595N | PFHB1A | S3IV |
| T1620K | PFHB1A | S4IV |
| T220I | SSS1 | S4I |
| A735V | SSS1 | S1II |
| P1298L | SSS1 | S3-4III |
| G1408R | SSS1 | SF III |
| D1792N | SSS1 | CTD |
| S1710L | VF1 | SF IV |
| F532C | SIDS | I-II linker |
| G1084S | SIDS | II-III linker |
| S1333Y | SIDS | S4-5III |
| F1705S | SIDS | SF IV |
| D1275N | ATRST1 | S3III |
| D1275N | CMD1E | S3III |
| M138I | ATFB10 | S1I |
| E428K | ATFB10 | S6I |
| H445D | ATFB10 | I-II linker |
| N470K | ATFB10 | I-II linker |
| A572D | ATFB10 | I-II linker |
| E655K | ATFB10 | I-II linker |
| T1131I | ATFB10 | II-III linker |
| R1826C | ATFB10 | CTD |
| V1951M | ATFB10 | CTD |
| N1987K | ATFB10 | CTD |

**PFHB1A**: Progressive familial heart block 1A; **SSS1**: Sick sinus syndrome; **VF1**: Familial paroxysmal ventricular fibrillation 1; **SIDS**: Sudden infant death syndrome; **ATRST1**: Atrial standstill 1; **CMD1E**: Cardiomyopathy, dilated 1E; **ATFB10**: Atrial fibrillation, familial, 10.

**Supplementary Table 4 | Activation and steady-state inactivation parameters of $Na_v1.5$ transiently co-expressed with or without β1 in HEK293T cells.**

| | Parameters | $Na_v1.5$ | $Na_v1.5$+β1 |
|---|---|---|---|
| **Activation** | $V_{1/2}$ (mV) | -36.76 ± 0.58 | -31.93 ± 0.42 |
| | P | / | 0.1593 |
| | slope | 6.95 ± 0.51 | 7.46 ± 0.37 |
| | P | / | 0.3808 |
| | n | 5 | 6 |
| **Inactivation** | $V_{1/2}$ (mV) | -74.77 ± 0.62 | -67.92 ± 0.80 |
| | P | / | 0.0511 |
| | slope | -11.70 ± 0.56 | -12.20 ± 0.76 |
| | P | / | 0.4879 |
| | $\tau_{inac}$ (ms) | 1.79 ± 0.32 | 2.35 ± 0.32 |
| | P | / | 0.2532 |
| | n | 5 | 5 |
| **Conductance** | $G_{top}$ (nS/pF) | 0.96 ± 0.07 | 1.24 ± 0.14 |
| | P | / | 0.1344 |
| | n | 5 | 6 |

Each data point represents mean ± s.e.m (standard deviation of mean) and *n* is the number of experimental cells from which recordings were obtained. Statistical significance was assessed using an unpaired t-test with Welch's correction.

**Supplementary Table 5 | Statistics for data collection and structural refinement.**

| | $Na_v1.5T$ | $Na_v1.5Q$ |
|---|---|---|
| **Data collection** | | |
| EM equipment | Titan Krios (Thermo Fisher Scientific Inc.) | |
| Voltage (kV) | 300 | |
| Detector | Gatan K2 Summit | |
| Energy filter | Gatan GIF Quantum, 20 eV slit | |
| Pixel size (Å) | 1.091 | |
| Electron dose ($e^-/Å^2$) | 48 | |
| Defocus range (μm) | -1.0 ~ -2.5 | |
| Number of collected micrographs | 5780 | 7799 |
| **Reconstruction** | | |
| Software | RELION 3.0 | |
| Number of used particles | 145,139 | 124,954 |
| Symmetry | C1 | |
| Resolution (Å) | **3.4** | **3.3** |
| Resolution for pore region (Å) | **3.3** | **3.2** |
| Map sharpening B-factor ($Å^2$) | -139.94 | -80 |
| **Refinement** | | |
| Software | Phenix | |
| Cell dimensions | | |
| a=b=c (Å) | 261.84 | 261.84 |
| α=β=γ (°) | 90 | 90 |
| Model composition | | |
| Protein residues | 1151 | 1151 |
| Side chains assigned | 1151 | 1151 |
| Sugar | 9 | 9 |
| R.m.s deviations | | |
| Bonds length (Å) | 0.01 | 0.01 |
| Bonds angle (°) | 0.69 | 0.68 |
| Ramachandran plot statistics (%) | | |
| Preferred | 90.92 | 92.14 |
| Allowed | 8.82 | 7.34 |
| Outlier | 0.26 | 0.52 |
